# Minimal medical imaging can accurately reconstruct geometric bone models for musculoskeletal models

**DOI:** 10.1101/432310

**Authors:** Edin K. Suwarganda, Laura E. Diamond, David J. Saxby, David G. Lloyd, A. Killen Bryce, Trevor N. Savage

**Author notes:** Current Address: School of Allied Health Sciences, Griffith University, Parklands Drive, Southport, Gold Coast, Queensland 4222, Australia (David G. Lloyd).

## Abstract

Accurate representation of subject-specific bone anatomy in lower-limb musculoskeletal models is important for human movement analyses and simulations. Mathematical methods can reconstruct geometric bone models using incomplete imaging of bone by morphing bone model templates, but the validity of these methods has not been fully explored. The purpose of this study was to determine the minimal imaging requirements for accurate reconstruction of geometric bone models. Complete geometric pelvis and femur models of 14 healthy adults were reconstructed from magnetic resonance imaging through segmentation. From each complete bone segmentation, three sets of incomplete segmentations (set 1 being the most incomplete) were created to test the effect of imaging incompleteness on reconstruction accuracy. Geometric bone models were reconstructed from complete sets, three incomplete sets, and two motion capture-based methods. Reconstructions from (in)complete sets were generated using statistical shape modelling, followed by host-mesh and local-mesh fitting through the Musculoskeletal Atlas Project Client. Reconstructions from motion capture-based methods used positional data from skin surface markers placed atop anatomic landmarks and estimated joint centre locations as target points for statistical shape modelling and linear scaling. Accuracy was evaluated with distance error (mm) and overlapping volume similarity (%) between complete bone segmentation and reconstructed bone models, and statistically compared using a repeated measure analysis of variance (p<0.05). Motion capture-based methods produced significantly higher distance error than reconstructions from (in)complete sets. Pelvis volume similarity reduced significantly with the level of incompleteness: complete set (92.70±1.92%), set 3 (85.41±1.99%), set 2 (81.22±3.03%), set 1 (62.30±6.17%), motion capture-based statistical shape modelling (41.18±9.54%), and motion capture-based linear scaling (26.80±7.19%). A similar trend was observed for femur volume similarity. Results indicate that imaging two relevant bone regions produces overlapping volume similarity > 80% compared to complete segmented bone models. These findings have implications for improving movement analysis and simulation with subject-specific musculoskeletal models.

## Introduction

Musculoskeletal (MSK) lower-limb models are ubiquitous tools used in motion analysis and simulation. Individual variation in bone anatomy influences muscle attachments (i.e. origins and insertions), and thus muscle-tendon-unit paths and moment arms. Consequently, individual variation in bone anatomy influences estimates of muscle-tendon-unit forces, joint contact forces [1,2], and articular mechanisms [3] in these models. Furthermore, combined variation in bone anatomy and muscle-tendon-unit force can influence finite element analysis of bone [4], cartilage [5,6], and tendon [7]. Therefore, generating accurate geometric bone models for subject-specific MSK models is important for human movement analysis and simulation.

Subject-specific geometric bone models can be reconstructed from 3-dimensional (3D) medical imaging through segmentation [3], or through mathematical methods that morph a template bone model to match individual bone geometry [8,9]. Medical image processing software is used to segment pixels corresponding to bones in images from x-ray computed tomography (CT) and magnetic resonance imaging (MRI), from which geometric bone models can be reconstructed. Currently, automated segmentation of bone is only possible with CT imaging [10], which, unlike high-fidelity MRI, cannot image soft tissues (i.e. muscle, ligament, cartilage, and tendon), which are valuable for MSK modelling and/or finite element analyses. However, segmentation of complete lower-limb bone models from MRI may take up to 11 hours [3]. Consequently, reconstructing geometric bone models through segmentation of MRI is resource intensive (i.e. cost and time), making it impractical for studies with large sample sizes or repeated measures.

As an alternative to complete segmentation from medical imaging, mathematical methods can be used to reconstruct geometric bone models from incomplete medical imaging segmentation of bone and/or anthropometric measurements by morphing a template bone model. Examples of mathematical morphing methods include linear scaling, free-form deformation (i.e. host-mesh fitting), parametrised nodal fitting (i.e. local-mesh fitting), and statistical shape modelling (SSM) [11,10,12]. Standard musculoskeletal modelling software, OpenSim [13], applies linear scaling to generic template bone models to personalise models to the individual. Linear scaling applies one to three orthogonal scale factors to template bone models to approximate individual bone dimensions. Scale factors are typically determined from motion capture (MOCAP) markers placed atop bony landmarks and/or using anthropometric measurements. However, MOCAP-based linear scaling results in pelvis and femur bone dimensions with width and/or depth errors of ∼5–20 mm compared to measurements from medical images [14]. Therefore, MOCAP-based linear scaling inadequately represent subject-specific bone geometry. Alternatively, mesh-fitting techniques (i.e. host-mesh and local-mesh fitting) morph template bone models (i.e. meshes) to individual bone geometry segmented from imaging data. Mesh-fitting is an iterative morphing processes that minimises distance error between segmentation and closest points on the template bone model. Host-mesh fitting applies free-form deformation to the template bone model, aiming to minimise distance error by deforming a host bounding box in which the template bone model is embedded [8]. Local-mesh fitting is a parametric fitting method, where individual nodes on the template bone model undergo translation to minimise distance error [15]. Mesh-fitting works with complete bone segmentation, but may not adequately adjust (e.g. collapse meshes, surface artefacts) regions of the template bone model unaccounted for in the incomplete medical imaging [8]. Thus, it remains unclear that these fitting methods alone may accurately reconstruct geometric bone models.

Data mining methods, such as SSM, analyse variation in bone anatomy within a training set and can be used to reconstruct geometric bone models. Reconstruction using SSM morphs bone shape variance along principal components (PCs) of variance, to minimise the distance error to bone segmentation from medical images and/or anthropometric measurements [11]. A SSM has been used previously to reconstruct bone models from incomplete medical imaging [16], and the resulting bone models had more accurate anatomic coordinate systems compared to linear scaling. Additionally, Nolte and colleagues [9] demonstrated geometric models of femur and tibia/fibula can be accurately (average distance error 0.50±0.33 mm) reconstructed from incomplete bone segmentations (two bone regions) compared to complete bone segmentation. Though SSM has accurately reconstructed geometric models of lower-limb long bones from incomplete bone segmentation, the minimal medical imaging required to accurately reconstruct geometric bone models is yet to be investigated.

The purpose of this study was to compare the accuracy of pelvis and femur geometric models reconstructed from different levels of imaging incompleteness to identify the minimal imaging requirements for accurate subject-specific bone models. We compared the reconstruction accuracy of pelvis and femur geometric models using target data comprising complete sets of bone segmentations, three incomplete sets of bone segmentations (each representing different levels of imaging incompleteness), and two MOCAP-based methods. We hypothesised that bone models reconstructed from more complete sets would be more accurate compared to those reconstructed from less complete sets and MOCAP-based methods.

## Methods

This cross-sectional study was approved by the institutional human research ethics committee. All participants provided written informed consent prior to testing. Fourteen healthy adults (age = 28.79±4.82 years, height = 1.73±0.85 m, mass = 69.17±16.22 kg), who reported no history of lower-limb injury and were free from cardiovascular and neuromuscular conditions, participated.

### Magnetic resonance imaging, bone segmentation, and motion capture

Complete pelvis and right-side femur bones were imaged using a Philips Ingenia 3 T MRI unit (Eindhoven, Netherlands). Image sequences were optimised for bone and muscle visibility, consisted of ∼1100 axial images, 560×560 pixel in-plane resolution, no inter-slice gap, and reconstructed voxel dimension 0.77 mm × 0.77 mm × 1.00 mm. Complete pelvis (excluding sacrum) and femur geometric models were reconstructed by segmenting pixels corresponding to the respective bones in each image slice using Mimics version 17 (Materialise, Leuven, Belgium).

From each complete set of bone segmentations, three sets of incomplete bone segmentations were created (Fig 1.). Incomplete bone sets consisted of proximal, middle, and/or distal bone regions, with each region truncated superiorly and inferiorly. Pelvis truncation was performed at the acetabula, ilia, and ischiopubic rami, while femur truncation was performed at proximal femur, epicondyles, and mid-shaft. For both pelvis and femur, incomplete segmentation set 1 was composed of one geometrically-complex bone region (i.e. femoral head or pelvic acetabulum), set 2 was composed of set 1 plus a second large bone region, and set 3 was composed of set 2 plus a third small bone region (Fig 1.). For the pelvis, set 1 consisted of the acetabula only, truncated superiorly and inferiorly at the acetabulum edge. Set 2 consisted of set 1 plus the ilia, truncated by parallel cuts emanating at the posterior superior and inferior iliac spines and terminating at the anterior superior iliac spine. Set 3 consisted of set 2 plus part of the ischiopubic rami. For the femur, set 1 consisted of femoral head and greater trochanter, truncated at ∼10 mm of proximal bone end and inferiorly at approximately half the distance between greater and lesser trochanters. Set 2 consisted of set 1 plus femoral epicondyles, truncated from the start of the femoral shaft and at ∼5 mm of distal bone end. Set 3 consisted of set 2 plus ∼10 mm of femoral shaft, located midway between femoral head and epicondyles. Complete and incomplete sets were used to reconstruct pelvis and femur models, and to evaluate geometric bone model accuracy in comparison to complete bone segmentations.

**Fig 1.**
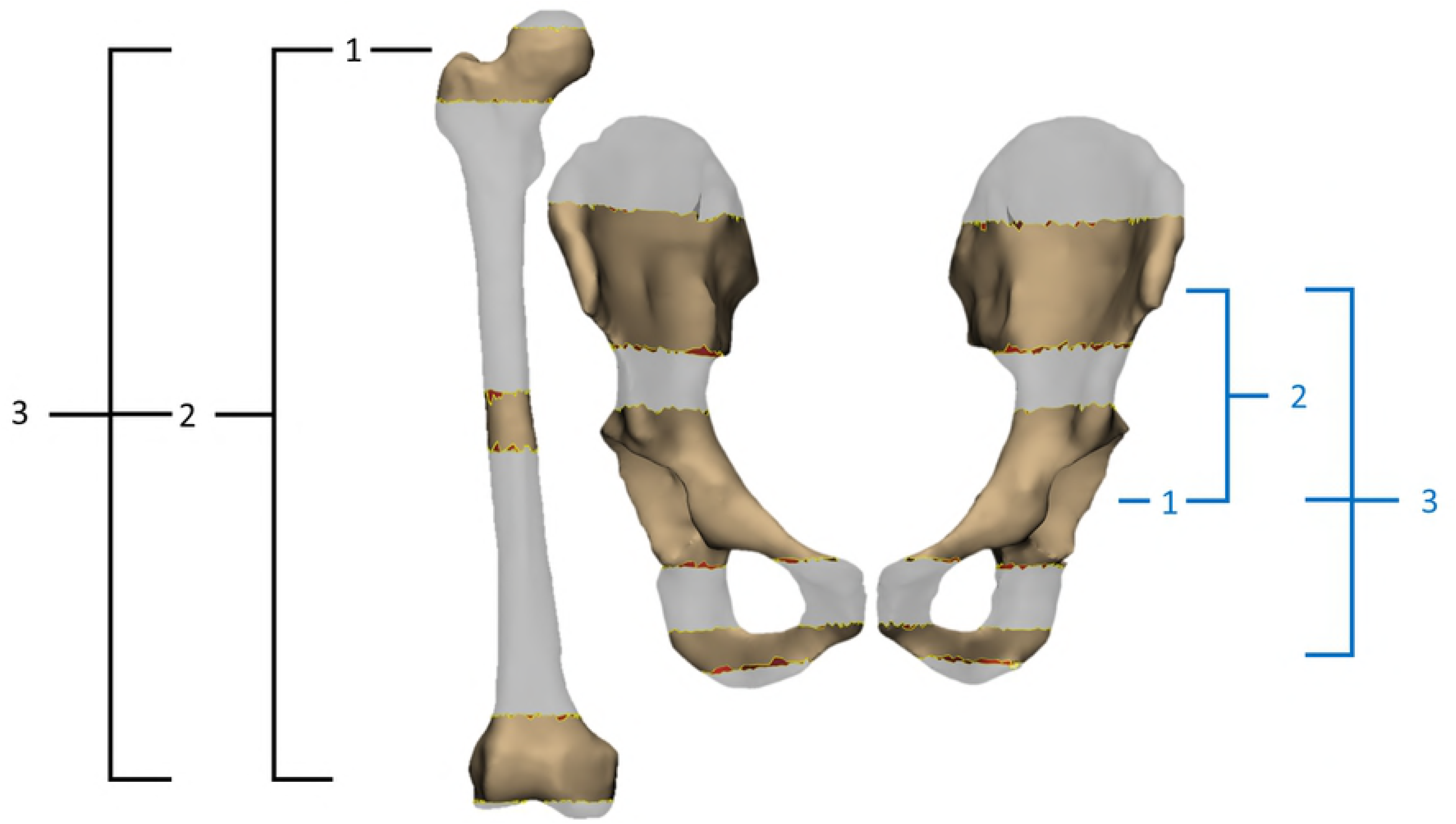
Example of proximal, middle, and distal bone regions for incomplete sets of the femur and pelvis created from complete bone segmentations from magnetic resonance imaging. Femur (black brackets) and pelvis (blue brackets) segmentation sets 1 through 3, with each bone region truncated superiorly and inferiorly. Incomplete pelvis sets were truncated at the acetabula, ilia, and ischiopubic rami, and femur truncation was performed at proximal femur, epicondyles, and mid-shaft.

Motion capture data were also used to reconstruct geometric bone models. Retro-reflective markers were placed on the skin surface atop bony landmarks while subjects remained in upright stance. Markers were placed according to the University of Western Australia marker set [17]. Instantaneous marker positions were acquired with Vicon Nexus version 1.8 at 100 Hz using a 12-camera MOCAP system (Vicon Motion Systems, Oxford, UK).

### Bone reconstruction

Morphing techniques were used to reconstruct geometric bone models using target data from complete and incomplete sets of pelvis and femur segmentations from MRIs or MOCAP marker data. Morphing was undertaken using the free and open-source software the Musculoskeletal Atlas Project (MAP) Client [18] or simple linear scaling of the OpenSim bones. Morphing via the MAP Client using the complete sets of pelvis and femur segmentations was used as the gold standard for geometric bone reconstructions.

The MAP Client employs SSM with PC, host-mesh and local-mesh morphing methods. The lower limb bones of the MAP Client SSM’s were trained using PC analysis performed on x-ray CT segmentations of cadaveric bones (214 femurs, and 26 full lower-limbs). The MAP Client SSM for each bone consists of a tessellation of higher-order (3^rd^ and 4^th^ order) Lagrange piece-wise elements with a fixed number of nodes and boundaries shared by adjacent elements (Fig 2). Focusing on SSM’s of the pelvis and femur, increasing the number of PCs increases the total variance accounted for in the resulting bone models and reduces distance errors, but also accounts for more individual variation from the training set (S1 Appendix). The optimal number of PCs (n = 4) for pelvis and femur SSM was established by minimising distance error, while using the least numbers of PCs [19] (S2 and S3 Appendix).

**Fig 2.**
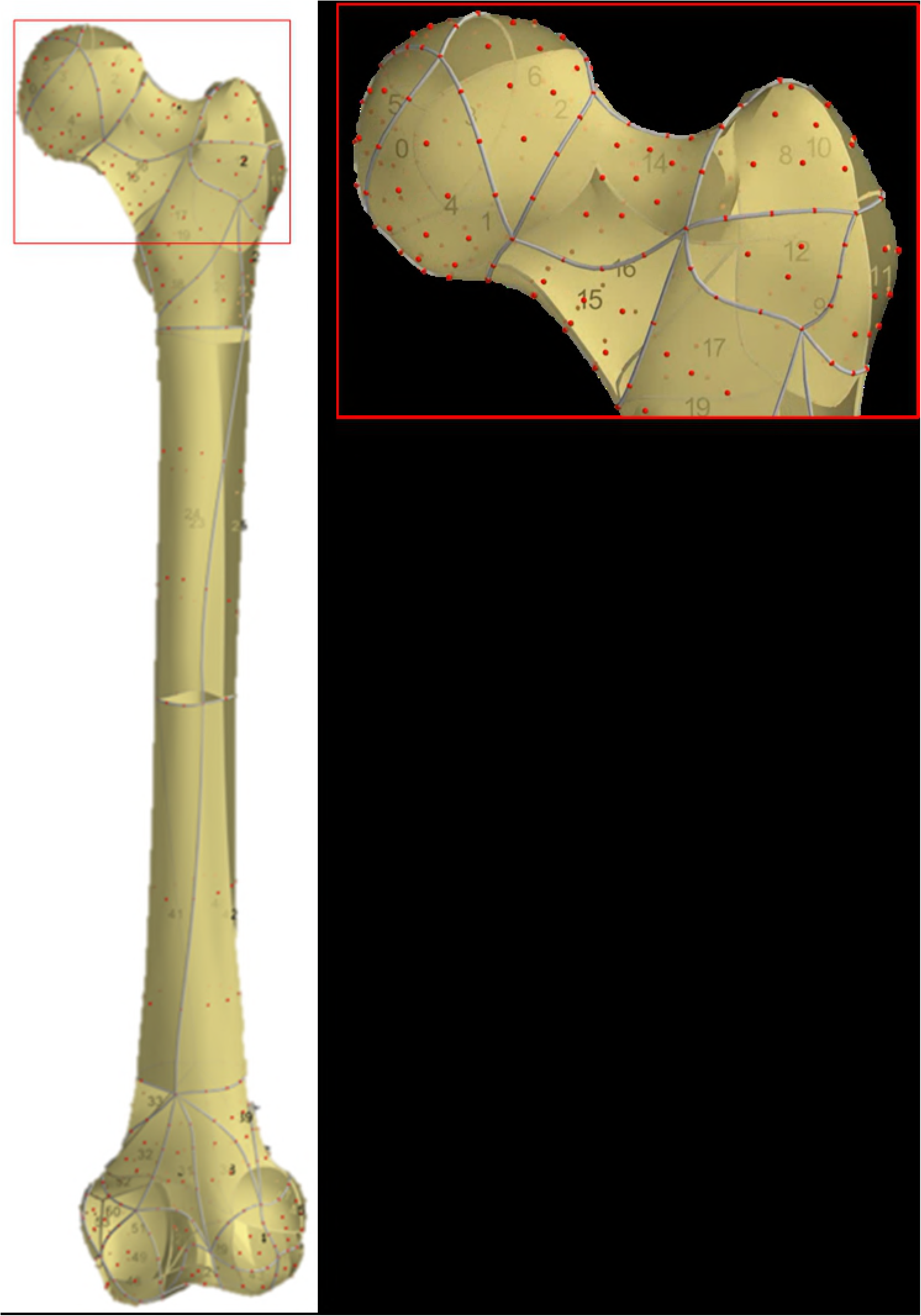
The Musculoskeletal Atlas Project Client statistical shape model for femur, which consists of Lagrange piece-wise elements and element nodes. Statistical shape model elements are 2-dimensional, higher-order, quadrilateral or triangular, and consists of fixed element numbering (yellow sections), fixed node distribution (red dots), and shared boundaries (grey lines) with adjacent elements. Figure inset – enlargement of proximal femur.

Prior to bone model reconstructions via the MAP Client, the average SSMs were rigidly registered to the bone segmentation sets to minimise distance error using an iterative closest-point method [20]. Subsequently, bone model reconstructions proceeded using SSM, followed by host-mesh fitting, and then local-mesh fitting (henceforth referred to as “MAP Client morphing process”). SSM was performed first since it was the coarsest morphing technique with the fewest degrees of fitting freedom (using 4 PCs). Local-mesh fitting was performed last, because it had the greatest degrees of fitting freedom, adjusting each individual mesh node (pelvis contains 1946 nodes and femur contains 634 nodes). Host-mesh fitting was performed second as it had intermediate degrees of fitting freedom. Finally, using a subset of participants (n = 5) we also showed that smaller distance errors resulted from combinations of morphing techniques (SSM, host-mesh, and local-mesh fitting) compared to SSM alone (S4 Appendix).

Each morphing technique (SSM, host-mesh and local-mesh fitting) was constrained using penalty weights to ensure bone surface smoothness and prevent unnatural shape. With SSM, penalty was applied to the Mahalanobis distance, which quantifies the similarity of morphed and mean SSM. The higher the Mahalanobis distance, the more dissimilar the morphed and mean SSM, and the penalty weight (range 0–1) was set to 0.1. Host-mesh fitting involved three different smoothing terms. The first pertained to the host bounding box and was a 3D second-order weighted Sobolev term [21]. The second and third terms pertained to the slave mesh (i.e. embedded SSM) and were a 2D second-order weighted Sobolev term and smoothing term based on piece-wise element boundary normal, respectively. The 2D Sobolev smoothing weights penalise high curvature within piece-wise elements (i.e. along boundaries and across surfaces) [22]. Boundary normal smoothing weights [15] penalise large angles between normals of nodes shared by adjacent piece-wise elements. Both Sobolev and boundary normal smoothing reduce occurrences of mesh self-intersection and creasing [13]. In local-mesh fitting, penalties were applied through the 2D Sobolev term and boundary normal smoothing terms. The 2D Sobolev penalties were larger for bone model reconstruction from incomplete (10e^-3^ to 10e^-5^) sets to constrain morphing at open end bone truncations relative to complete (10e^-5^ to 10e^-6^) bone segmentations.

Bone models reconstructed from MOCAP data used 3D marker positions to identify surface landmarks and joint centre locations as target points (Fig 3). Hip joint centre locations were estimated from marker positions using the equations of Harrington and colleagues [23], while knee joint centres were estimated to be the average of medial and lateral femoral condyles. For the MAP Client, SSM was only used with 4 PCs to fit nodal points to MOCAP target points [24]. The same MOCAP target points were also used to linearly scale generic bone models from OpenSim version 3.3 [13] using a custom Python program (Python Software Foundation, Delaware, United States). Scaling factors for each bone’s orthogonal dimensions were determined by the quotient of the distance between two (or more) MOCAP makers defining bony landmarks or joint centre locations with corresponding virtual markers on the generic bone models.

**Fig 3.**
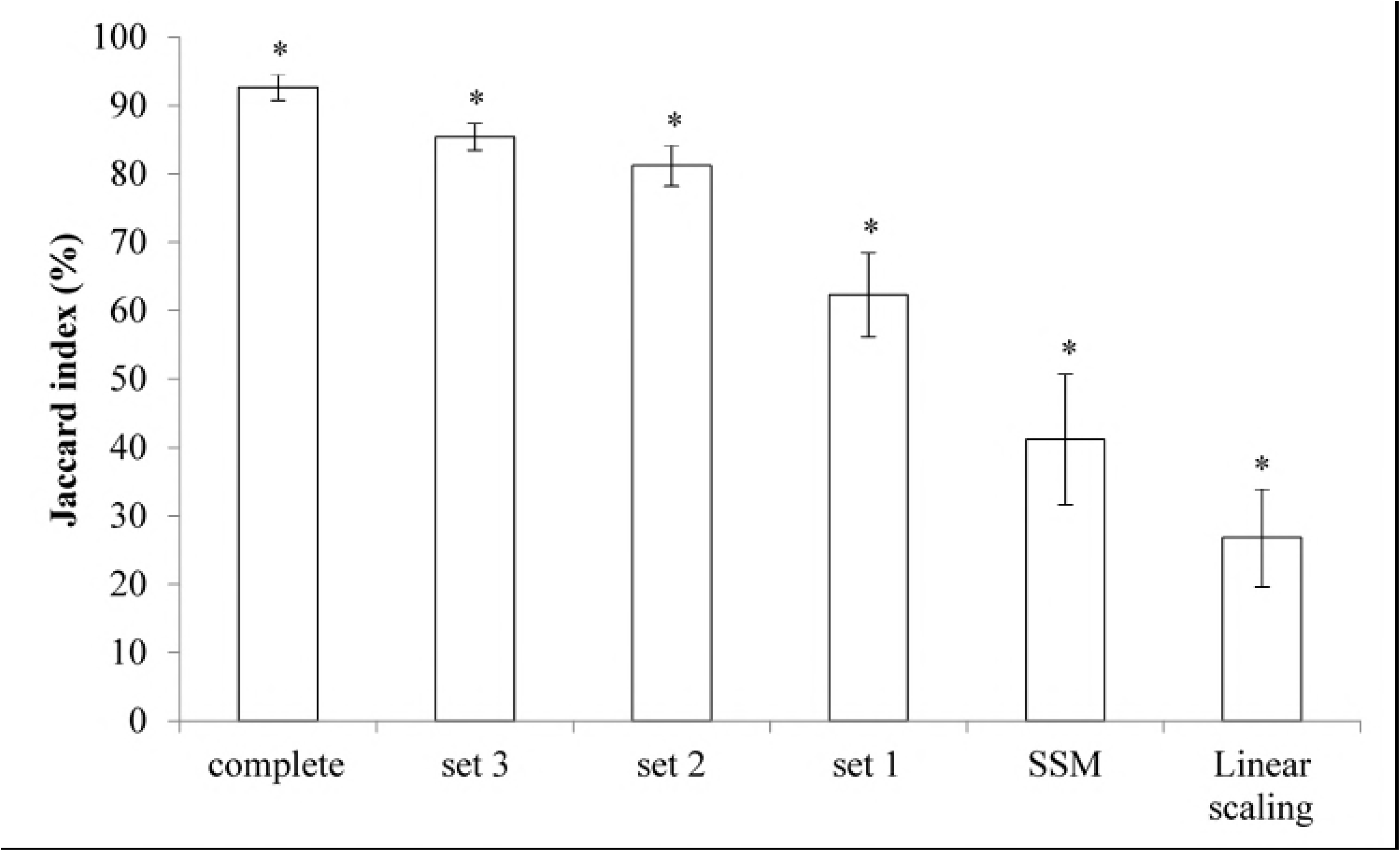
Overview of subject-specific bone reconstruction methods performed using both magnetic resonance imaging (MRI) and motion capture (MOCAP) data. Four different MRI segmentation sets were used: one complete and three sets of incomplete bone segmentations were created for pelvis and femur for each subject to reconstruct subject-specific geometric bone models through the MAP Client morphing process (i.e. statistical shape modelling (SSM), followed by host-mesh, and then local-mesh fitting). Motion capture marker positions on pelvis and femur bony landmarks were used to calculate pelvis dimensions, as well as hip joint and knee joint centres. These landmarks and joint centres were then used to 1) perform SSM and 2) calculate three orthogonal scaling factors for each bone type for linear scaling of generic OpenSim bone models. Then, bone reconstructions from MOCAP-based methods were rigidly registered to the complete bone segmentation. In total, six bone models were generated for each bone type (excluding pelvis sacrum) and each subject, and subsequently evaluated for modelling accuracy using root mean square error (RMSE) and Jaccard index.

### Analyses

Reconstruction accuracy of pelvis and femur geometric models were evaluated using two metrics. First, the distance error between complete bone segmentation (i.e. a point cloud of the entire bone) and the closest nodes on reconstructed bone models was calculated using the root mean square error (RMSE, mm). The RMSE is an average error which has been applied in previous studies for comparing bone reconstruction accuracy [9, 16, 24]. Second, the overlapping volume similarity of complete bone segmentation and reconstructed geometric bone model was calculated using the Jaccard index (%) [25]. The Jaccard index was calculated in addition to the RMSE, because it accounts for absolute differences in overlapping volume of geometric bone models. The accuracy metrics of the different bone reconstruction methods were statistically compared (p<0.05) using a repeated measure analysis of variance followed by multiple pairwise comparisons with Bonferroni adjustment in the Statistical Package for the Social Sciences version 25 (SPSS Inc., Chicago, United States).

## Results

Distance errors of geometric bone models reconstructed from (in)complete sets of bone segmentation through the MAP Client morphing process were significantly lower than MOCAP-based reconstructions of both pelvis and femur (Table 2). The distance error of reconstructed bone models from MOCAP-based SSM was also significantly lower than MOCAP-based linear scaling. Notably, femur models reconstructed from complete sets of segmentation had significantly higher distance error compared to reconstructions from incomplete sets 3.

**Table 2.**
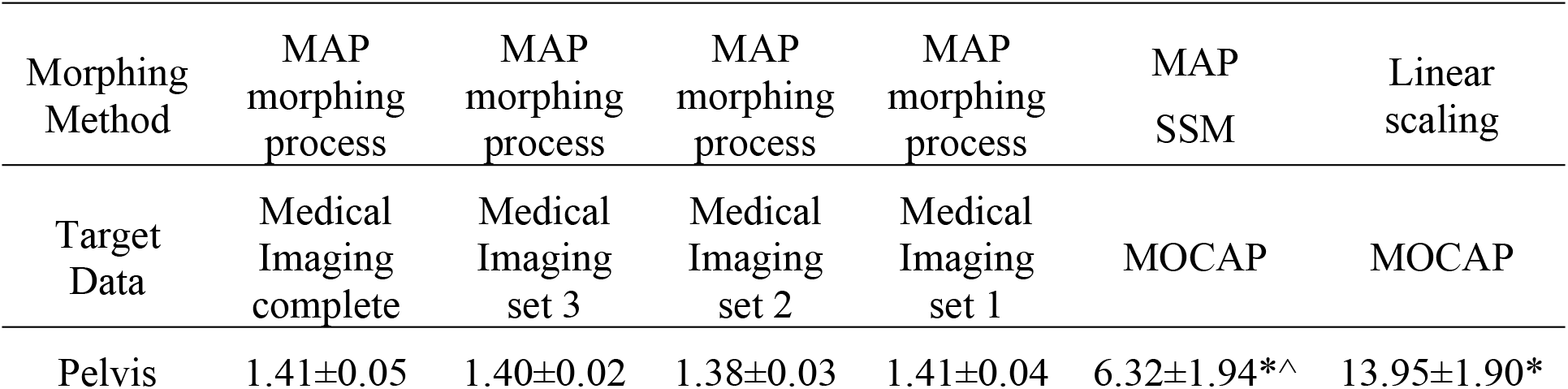
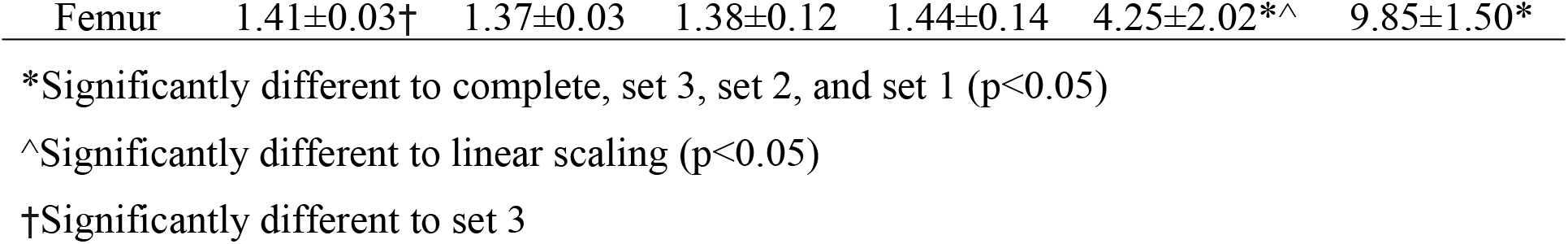
Comparison of root mean square error (RMSE, mm) for pelvis and femur bone models reconstructed by the different methods. Pelvis and femur geometric models were reconstructed from complete sets of bone segmentations, three incomplete sets of bone segmentations (sets 1 through 3, as depicted in Fig 1), motion capture (MOCAP)-based statistical shape modelling (SSM), and MOCAP-based linear scaling.

| Morphing Method | MAP morphing process | MAP morphing process | MAP morphing process | MAP morphing process | MAP SSM | Linear scaling |
| --- | --- | --- | --- | --- | --- | --- |
| Target Data | Medical Imaging complete | Medical Imaging set 3 | Medical Imaging set 2 | Medical Imaging set 1 | MOCAP | MOCAP |
| Pelvis | 1.41±0.05 | 1.40±0.02 | 1.38±0.03 | 1.41±0.04 | 6.32±1.94*<sup>^</sup> | 13.95±1.90* |
| Femur | 1.41±0.03† | 1.37±0.03 | 1.38±0.12 | 1.44±0.14 | 4.25±2.02*^ | 9.85±1.50* |

\*Significantly different to complete, set 3, set 2, and set 1 (p<0.05)
^Significantly different to linear scaling (p<0.05)
†Significantly different to set 3

Overlapping volume similarity for the pelvis bone models significantly reduced with the level of incompleteness. Jaccard indices were highest for pelvis geometric models reconstructed from complete sets (92.70±1.92%), reducing to 85.41±1.99%, 81.22±3.03%, 62.30±6.17% for incomplete sets 3, 2, and 1, respectively. Notably, Jaccard indices reduced significantly further in pelvis bone models reconstructed from MOCAP-based SSM (41.18±9.54%) and MOCAP-based linear scaling (26.80±7.19%) (Fig 4). A similar pattern of reduction in volume similarity was found for femur geometric models, except between reconstructions from incomplete sets 1 and MOCAP-based SSM (Fig 5). Jaccard indices of reconstructed femur bone models were 96.49±7.19%, 90.44±2.35%, 85.41±5.03%, 66.02±10.83%, 68.22±11.33%, and 49.38±11.57% for complete set, incomplete sets 3, 2, 1, MOCAP-based SSM, and MOCAP-based linear scaling, respectively.

**Fig 4.**
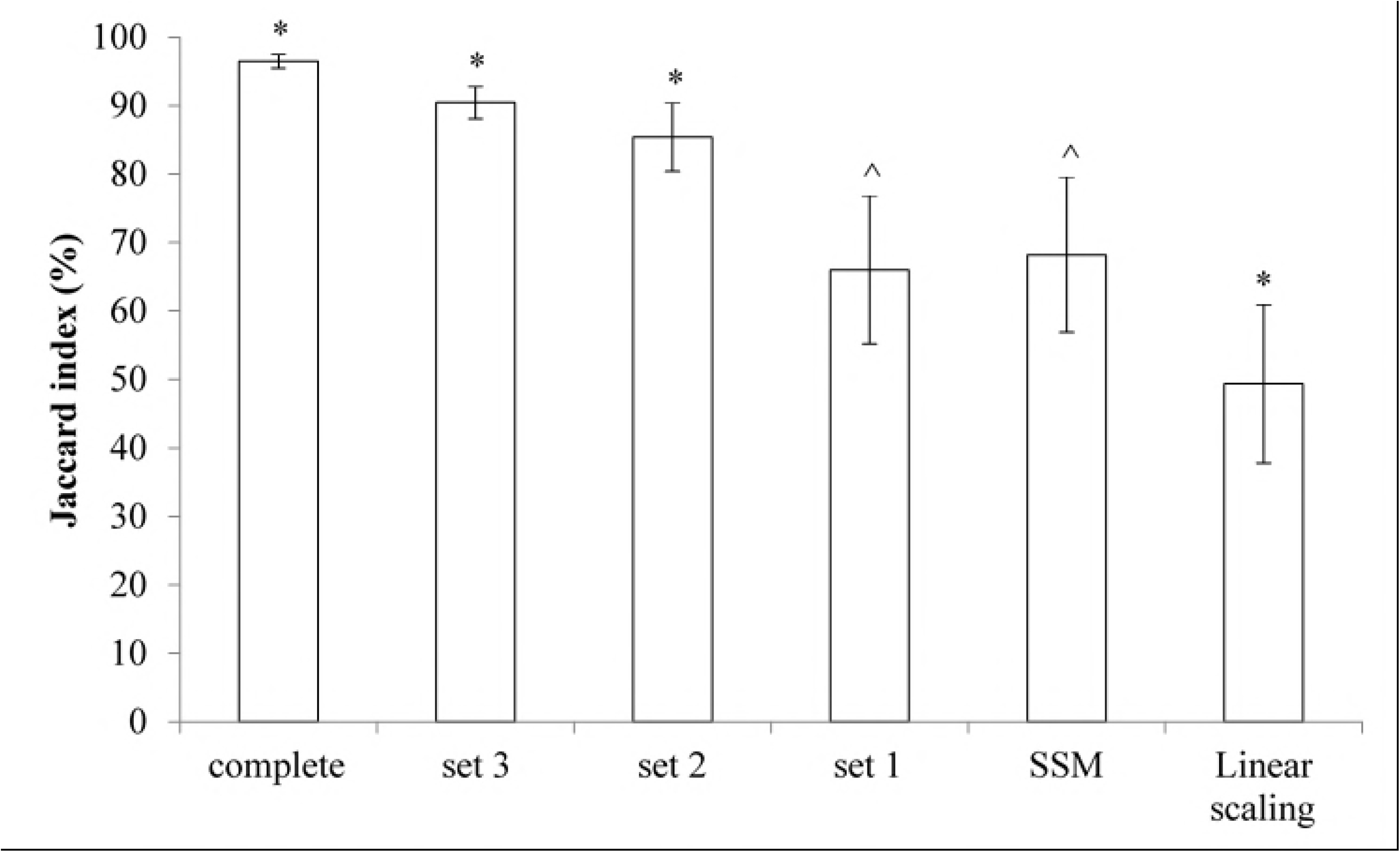
Comparisons of Jaccard index (%) for pelvis bone models reconstructed by the different methods. Pelvis geometric models were reconstructed from complete sets of bone segmentations, three incomplete sets of bone segmentations (sets 1 through 3, as depicted in Fig 1), motion capture (MOCAP)-based statistical shape modelling (SSM), and MOCAP-based linear scaling. Jaccard index (%), a measure for overlapping volume similarity, was calculated between complete bone segmentation and each reconstructed geometric pelvis model. Significant difference (*) from all reconstruction methods (p<0.05).

**Fig 5.**
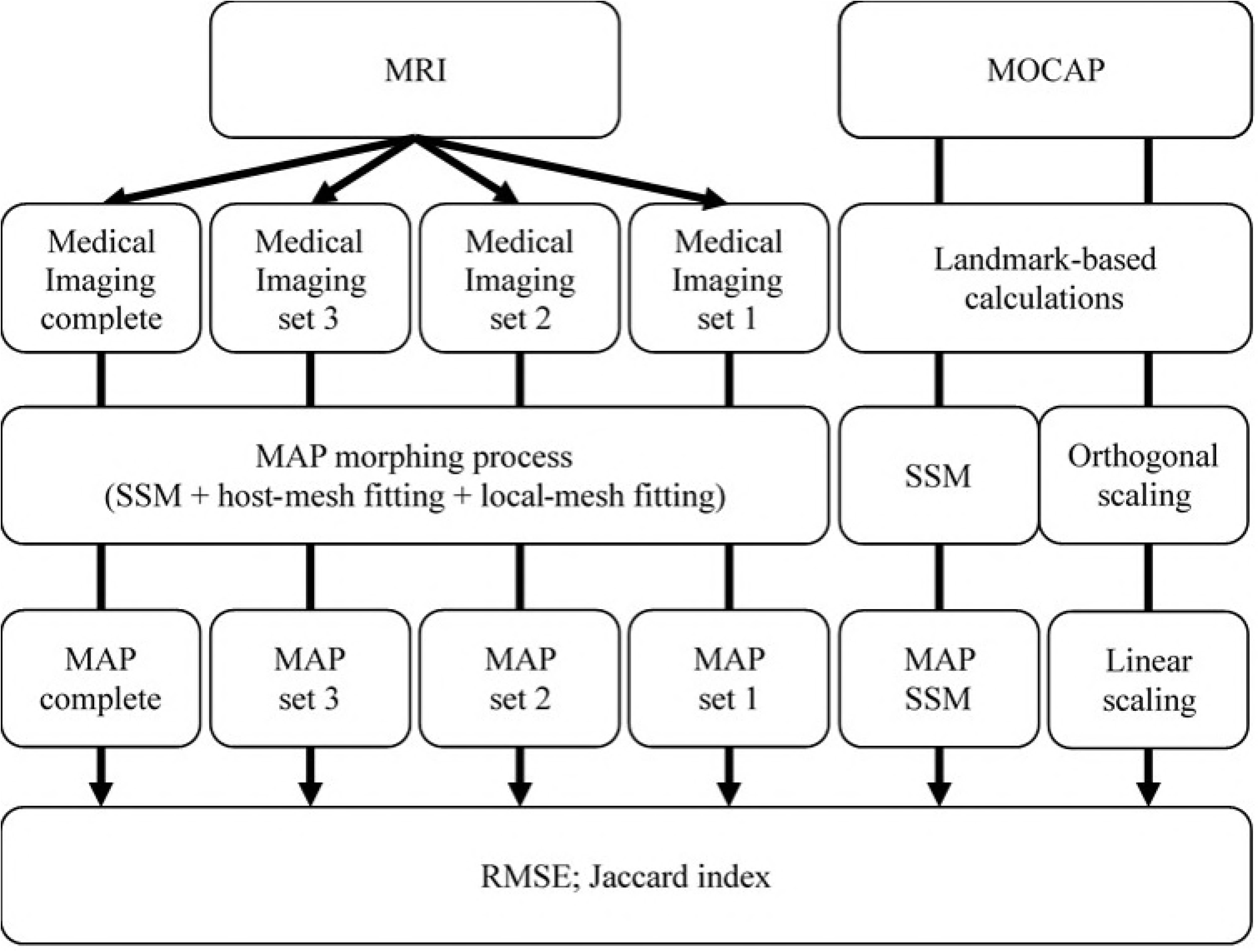
Comparisons of Jaccard index (%) for femur bone models reconstructed by the different methods. Femur geometric models were reconstructed from complete sets of bone segmentations, three incomplete sets of bone segmentations (sets 1 through 3, as depicted in Fig 1), motion capture (MOCAP)-based statistical shape modelling (SSM), and MOCAP-based linear scaling. Jaccard index (%), a measure for overlapping volume similarity, was calculated between complete bone segmentation and each reconstructed geometric femur model. Significant difference (*) from all reconstruction methods (p<0.05). Significant difference (^) from complete set, set 3, sets 2, and motion capture-based linear scaling (p<0.05).

## Discussion

The purpose of our study was to compare the accuracy of bone geometric models mathematically reconstructed from different levels of imaging incompleteness, and to identify the minimal imaging requirements for accurate representation of subject-specific bones. Our hypothesis, that geometric bone models reconstructed from more complete segmentations are more accurate than those reconstructed from less complete or MOCAP-based data, was partially supported. We found that pelvis and femur bone models reconstructed from different levels of imaging incompleteness had significantly different volume similarities compared to complete bone segmentations, but not significant different distance errors. Results suggest that imaging of only 2 bone regions, as represented by set 2 in this study, is sufficient (>80% volume similarity) to accurately reconstruct subject-specific pelvis and femur bone models.

Contrary to our hypothesis, pelvis and femur bone models reconstructed from different levels of incompleteness had comparable distance errors, indicating the MAP Client morphing process reconstructed accurate geometric models, even when reconstructed from incomplete sets of bone segmentations. Surprisingly, distance error of femur bone reconstructions from complete segmentation was significantly higher than reconstructions from incomplete set 3, which may be due to a low number of nodes on the femur shaft of the SSM (Fig 2). A low number of nodes on the femur shaft may have artificially raised the RMSE calculated between the large numbers of points on the femur shaft and closest points on the SSM. Given that RMSE, possibly may not capture meaningful geometric differences, pelvis and femur reconstructions from different levels of incompleteness show variations in overlapping volume similarity (Jaccard Index), which reveal a different interpretation of reconstruction accuracy.

Consistent with our hypothesis, pelvis geometric models had significant lower volume similarity (i.e. lower reconstruction accuracy) with the greater level of incompleteness, from complete sets, to incomplete sets 3, 2, and 1, MOCAP-based SSM, and linear scaling, respectively. Volume similarity of femur bone models showed a similar trend of significant reduction, except between femur reconstructions from incomplete set 1 and MOCAP-based SSM. Notably, higher volume similarity of reconstructions from incomplete sets is likely achievable when proximal and distal bone ends are imaged, as these constrain morphing that is meaningful to physical boundaries. Moreover, inclusion of bone terminals allows for a reduction in smoothing penalties, which has direct implications for reconstruction accuracy. Differences in volume similarity are likely caused by the differences in geometric bone regions used as inputs, and, to a lesser extent, from the different smoothing penalties applied to incomplete and complete sets of segmentation in the morphing process. Imposition of smoothing penalties caused overlapping volume similarity to be less than 100% for pelvis (∼93%) and femur (∼96%) geometric models reconstructed from complete segmentations. Although the application of smoothing constraints is necessary to prevent bone surface artefacts in regions of the template bone model unaccounted for in incomplete imaging it does produce imperfect reconstruction. Regardless of the need of smoothing constraints, volume similarity could distinguish reconstruction accuracy from different levels of imaging incompleteness.

Importantly, bone models reconstructed from incomplete set 2 had volume similarities of ∼80% and ∼85% for pelvis and femur, respectively. Reconstructions from complete sets improved volume similarity by ∼11% for both pelvis and femur compared to set 2 reconstructions. The practical relevance of improvements this size is unclear and depends, at least in part, on the impact downstream in MSK modelling and/or finite element analyses. This is an open research question and worthy of further investigation. Nonetheless, our findings relating to volume similarity suggest that an incomplete segmentation (i.e. set 2) is sufficient to reconstruct accurate (>80%) subject-specific geometric pelvis and femur bone models through the MAP Client morphing process, which is a marked improvement over bone models reconstructed through standard MOCAP-based linear scaling of generic bone models (i.e. +54.42% for pelvis and +36.04% for femur).

Consistent with previous findings [9, 16, 24] regarding the distance errors of reconstructions, our study has shown that the volume similarity of pelvis and femur bone models reconstructed through MOCAP-based SSM is significantly larger (+14.38% for pelvis and +18.84% for femur) than MOCAP-based linear scaling. The volume similarity of femur geometric models reconstructed from incomplete set 1 and MOCAP-based SSM were not significantly different, which may be explained by the least complete bone target data (i.e. single bone region and a set of anatomical target points) being used, the lower natural variance and/or complexity of femur bone shape in comparison to pelvis bone shape, and/or by the high percentage of variance (∼98%) accounted for by the femur SSM (using 4 PCs). Regardless of the reason, our findings corroborate previous investigations that show geometric bone model reconstruction via shape variance of SSM is more accurate compared to linear scaling.

With their custom-derived SSM, Nolte and colleagues [9] reconstructed femur geometric models from complete segmentations with RMSE lower than our result (∼0.50 mm and ∼1.41 mm respectively). A possible explanation for our higher RMSE is the nodal density, as their SSM was composed of > 10,000 nodes, whereas the MAP Client femur SSM consisted of only ∼600 nodes. When nodes from the SSM are fewer in number than the complete bone segmentation (multiples of ten thousand points when segmented from high resolution MRI), a single SSM node could serve as closest point for multiple segmentation points. Thus, a lower distribution of nodes will artificially increase the RMSE between closest points. The MAP Client SSM uses element-based fixed surface node numbering for three important reasons. First, to improve computation time of mathematical morphing. Second, to regulate bone smoothness and Lagrange continuity of piece-wise elements nodes, which is beneficial for subsequent finite element modelling. Third, to enable surface node allocation of muscle origin and insertion areas, muscle wrapping surfaces, and, implicitly, muscle pathways. This eliminates fitting of muscle origin and insertions and muscle pathway to resulting geometric bone models as proposed by Nolte and colleagues [9]. Nonetheless, both our approach and that of Nolte and colleagues [9] achieved RMSE within the distance error of adjacent imaging voxels (up to 2 mm), which means that further improvement on final RMSE is unnecessary. However, RMSE is sensitive to nodal distribution which differs across custom-derived SSM, and may suggest that distance error is less meaningful than volume similarity for the evaluation of geometric bone models.

This study has limitations that need to be considered. First, this study did not include all lower-limb bones, such as the patella, tibia/fibula bones, and bones comprising the feet. We chose to reconstruct geometric pelvis and femur models, because femur RMSE results could be evaluated against findings from the literature [9], and to evaluate similarity of femur model reconstruction accuracy to that of the pelvis (for which no findings from medical imaging based reconstruction is reported in the literature). Second, the nodal density of MAP Client SSM can potentially be increased by interpolation between fixed nodal points after the morphing process, which may alter RMSE calculation of the reconstructed geometric bone models. However, the nodal density of MAP Client SSM with which RMSE calculation would converge is unclear. Third, the reconstruction accuracy achieved through other combinations of morphing techniques is unknown beyond our preliminary evaluation of 5 participants. However, the diverse anthropometric range of this subset could suggest that differences between combinations of morphing techniques could remain similar. Fourth, mathematical techniques established for morphing and smoothing are specific to the MAP Client and may only be transferrable to other SSM frameworks by adapting the algorithms. Fifth, the population examined were healthy adults, and the methods and results of this study may change when considering populations with different physical characteristics (e.g. paediatric or pathologic). Importantly, future research should be directed to identifying optimal parameterisation of SSM for reconstruction of bone models including training set size, node distribution, and smoothing constraints.

## Conclusions

Volume similarity of reconstructed geometric pelvis and femur models reduces as imaging becomes more incomplete. Using the MAP Client morphing process (i.e. SSM, followed by host-mesh, and then local-mesh fitting), medical imaging of only 2 relevant and truncated bone regions (set 2) is sufficient to accurately reconstruct both pelvis and femur bone models of healthy adults. In the absence of MRI, MOCAP-based SSM in the MAP Client is superior to MOCAP-based linear scaling of pelvis and femur bone models. These findings may have important implications for improving analyses and simulation of adult subject-specific MSK models.

## Acknowledgements

The authors are grateful for support of Ben Kennedy for the collection of high quality MRI sequences at QScan branch in Southport, Australia and the support of Jillian Eyles (University of Sydney, Australia), Kim L. Bennell (The University of Melbourne, Australia), Libby Spiers (The University of Melbourne, Australia), Ju Zhang (University of Auckland, New Zealand), and Thor F. Besier (University of Auckland, New Zealand) for the concept of bone modelling from incomplete input.

## Supporting information

## S1 Appendix.

**Individual and cumulative percentage of bone shape variance explained by each principal component for the pelvis (n = 26) and femur (n = 214) included in the training set**.

## S2 Appendix. The optimal number of principal components for pelvis statistical shape modelling

The average optimal number of principal components (red circles projected on the x-axis), for pelvis reconstruction of each participant included of the subset (n = 5) was 4.2±1.1. Optimal numbers were calculated by minimising the distance error (RMSE) and the least number of principal components.

## S3 Appendix. The optimal number of principal components for femur statistical shape modelling

The average optimal number of principal components (red circles red circles projected on the x-axis), for femur reconstruction of each participant of the subset (n = 5) was 4.2±2.3. Optimal numbers were calculated by minimising the distance error (RMSE) and the least number of principal components.

## S4 Appendix. The distance error of pelvis and femur reconstructions from complete bone segmentation through different morphing processes

Distance error (RMSE, mm) (mean ± standard deviation) of pelvis and femur reconstructions from complete bone segmentation for each participants of the subset (n = 5) through statistical shape modelling (SSM) and combinations of/with host-mesh and local-mesh fitting techniques. Differences (p<0.05) were investigated using repeated measures analysis of variance and selected pairwise comparisons with Bonferroni adjustment.

